# The MERS-CoV receptor gene is among COVID-19 risk factors inherited from Neandertals

**DOI:** 10.1101/2020.12.11.422139

**Authors:** Hugo Zeberg, Svante Pääbo

## Abstract

In the current SARS-CoV-2 pandemic, two genetic regions derived from Neandertals have been shown to increase and decrease, respectively, the risk of falling severely ill upon infection. Here, we show that 2-8% of people in Eurasia carry a variant promoter region of the *DPP4* gene inherited from Neandertals. This gene encodes an enzyme that serves as a receptor for the coronavirus MERS-CoV and is currently not believed to be a receptor for SARS-CoV-2. However, the Neandertal *DPP4* variant doubles the risk to become critically ill in COVID-19.

## Main text

It was recently shown that the major genetic risk factor for falling severely ill when infected with the novel SARS-CoV-2 virus is inherited from Neandertals (Zeberg and Pääbo, 2020a). Furthermore, a protective genetic variant for severe COVID-19 located on chromosome 12 has been shown to be inherited from Neandertals (Zeberg and Pääbo, 2020b). In light of these findings, Neandertals haplotypes might be of particular importance in attempts to reveal the genetic factors that predispose individuals to severe COVID-19.

Early in the pandemic, the membrane-bound enzyme ACE2 was identified as a receptor for SARS-CoV-2 (Hoffmann *et al*. 2020). ACE2 has previously been shown to be a functional receptor for other coronaviruses, including HCoV-NL63 and SARS-CoV-1. The facts that males are more at risk for severe COVID-19 and that *ACE2* is located on the X chromosome suggest that the *ACE2* would emerge as a risk locus for severe COVID-19 in genetic association studies. However, to date we are not aware of that any association between *ACE2* and severity of COVID-19 have been described. MERS-CoV, a coronavirus that was discovered in 2012 and have caused outbreaks in several countries (de Groot *et al*. 2013), utilizes DPP4 as a receptor (Wang *et al*. 2013; Stalin Raj *et al*. 2013). DPP4 is a membrane-bound enzyme that degrades a number of biologically active peptides. It is involved in several physiological systems, including the regulation of glucose metabolism (Mentlein, 1999). Inhibitors of DPP4 are used to treat diabetes and have been suggested to affect COVID-19 outcomes in preliminary studies (reviewed in Lim *et al*. 2021).

We looked for single nucleotide polymorphisms (SNPs) that carry Neandertal-like alleles at the *ACE2* and *DPP4* loci in the 1000 Genomes Project (Auton *et al*. 2015) (Supplementary Material). In a region encompassing the entire *ACE2* gene and 50,000 base pairs up- and down-stream (chrX:15529156-15670192, *hg19*) we find two such SNPs. However, these SNPs are not in linkage disequilibrium (LD) (r^2^ = 4e-4) and thus do not form a contiguous haplotype. For *DPP4,* we find 39 such SNPs in the corresponding region (chr2:162798751-162981052, *hg19*). They form several haplotypes (r^2^>0.9) in and around the *DPP4* gene (Fig. 1A).

**Figure 1.**
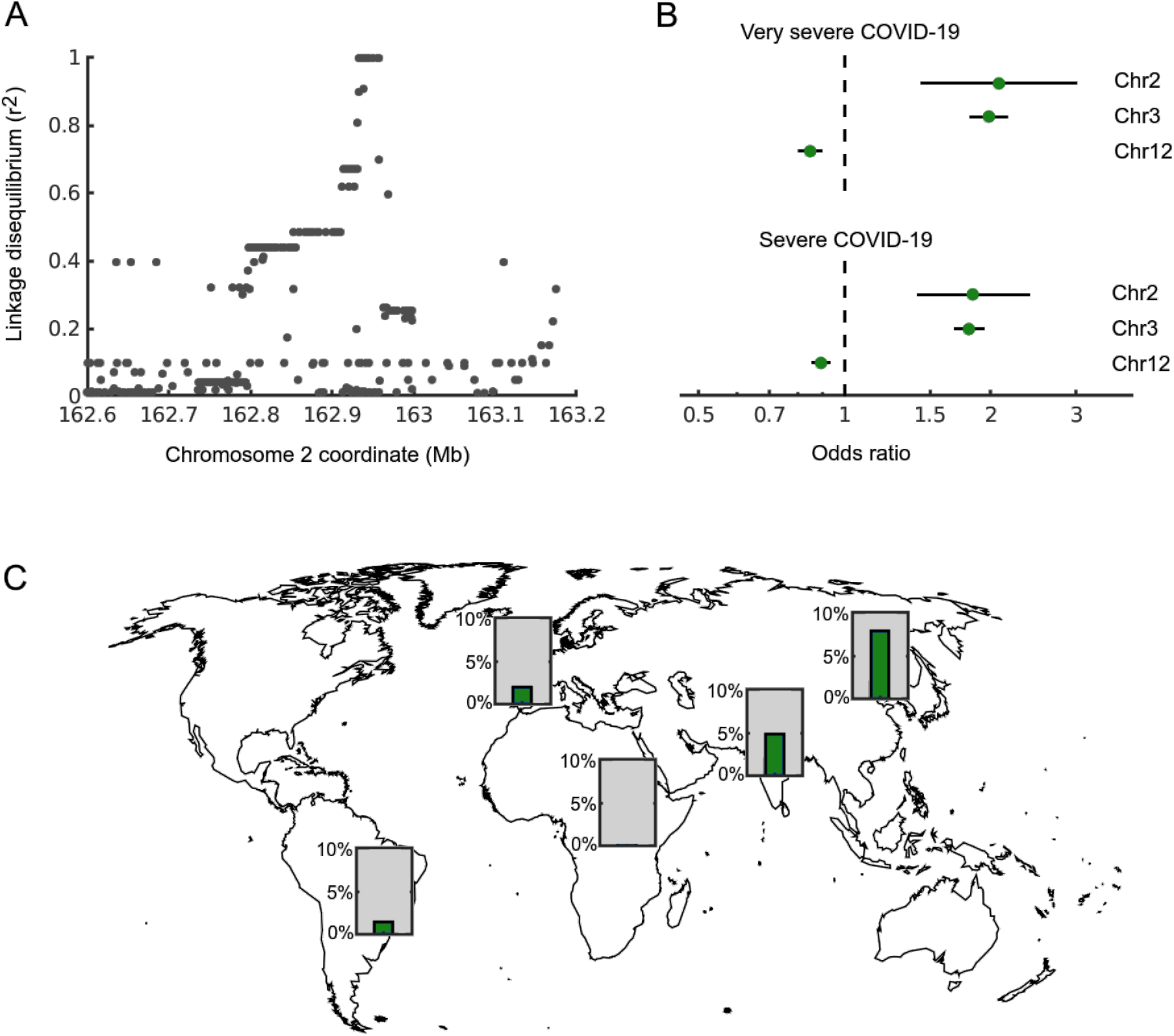
The Neandertal *DPP4* locus. **A)** Linkage disequilibrium to the index variant rs117460501 in Europeans in 1000 Genomes data (Auton *et al*. 2015). Note the step-wise decay in linkage disequilibrium, representing recombination events of longer Neandertal haplotypes. The core haplotype (r^2^ >0.8) covers the extended promoter of *DPP4* and has a length of 26.3 kb. **B)** Phenotypic impact of three Neandertal haplotypes on COVID-19 severity. The *DPP4* haplotype is located on chromosome 2, whereas the previously described risk and protective haplotypes are situated on chromosomes 3 and 12, respectively. Severe diseases is defined as an infection requiring hospitalization, whereas very severe disease requires active respiratory support beyond supplemental oxygen therapy. **C**) Carrier frequencies of the *DPP4* risk haplotype in continental populations. Data from the 1000 Genomes Project.

To investigate if the Neandertal haplotypes in *DPP4* are associated with severe COVID-19 we used the latest release of the COVID-19 Host Genetics initiative (HGI) (The COVID-19 HGI, 2020). We find that under the rare disease assumption, the Neandertal-like alleles are associated with ~80% increased risk per allele of being hospitalized upon infection with SARS-CoV-2 (Supplementary Table S1). The most strongly associated SNP (rs117888248) has an odds ratio of 1.84 (95% CI: 1.41-2.41, p = 7.7e-6). The risk for carriers of this allele of requiring mechanical ventilation is increased by ~109% (OR = 2.09, 95% CI: 1.44-3.03, p = 1.2e-4). The Neandertal-like alleles form a ~26.3 kb-haplotype (r^2^>0.8). Of the 15 SNP defining the haplotype (Supplementary Table S1), 14 carry alleles seen in hetero- or homozygous forms in a Neandertal genome (Prüfer *et al.* 2017). This haplotype is derived from Neandertals (p = 0.023) according to a published formula (Huerta-Sanchez *et al.* 2014) and using parameters as previously described (Zeberg and Pääbo, 2020a) (Supplementary Materials). It covers the extended promoter region of *DPP4.*

Thus, the risk of becoming severely ill in COVID-19 when carrying the Neandertal haplotype at the *DPP4* locus is of similar magnitude as the ~100% increase in risk associated with the previously described Neandertal haplotype on chromosome 3 (Zeberg and Pääbo 2020a) (Fig. 1B). Both these risk haplotypes have stronger effect sizes than the protective Neandertal haplotype on chromosome 12, which decreases the risk of becoming severely ill by ~23% (Zeberg and Pääbo 2020b). The Neandertal *DPP4* haplotype is present in ~1% of Europeans, ~2.5% in South Asians ~4% in East Asians, and ~0.7% in admixed Americans (Fig. 1C). It is absent among Africans south of the Sahara.

We calculated the statistical significance of the association between the Neandertal *DPP4* haplotype and severe COVID-19 under the null-hypothesis that Neandertal haplotypes have no impact on COVID-19. Because only a fraction of the Neandertal genome is found among present-day humans, and because Neandertal haplotypes are on average longer than other haplotypes, the statistical power to detect associations with Neandertal haplotypes is better than for genome-wide analyses. When we consider Neandertal haplotypes that are present in a frequency of >1% among Europeans in the 1000 Genomes Project and are identified in previously published maps of Neandertal contributions, we find that the effective number of hypotheses is 5,761, yielding an ‘introgression wide’ significance threshold of 8.7e-6 (Supplementary Material). Thus, under the null-hypothesis that Neandertal gene variants has no impact on COVID-19, the association of the *DPP4* haplotype with severe disease is significant.

It was recently shown that the spike protein of SARS-CoV-2 binds to DPP4 (Li *et al*. 2020) and that DPP4 is a functional receptor for HIV (Li *et al.* 2020). Moreover, *DPP9*, a homolog of *DPP4*, is associated with severe COVID-19 (Pairo-Castineira *et al*. 2020). However, one report suggest that SARS-CoV-2 does not use DPP4 as a receptor (Zhou *et al.* 2020). Nevertheless, the current findings suggest that the interaction of SARS-CoV-2 with membrane-bound and secreted forms of DPP4 deserves further investigation. Inhibitors of DPP4, which can be administered orally and are used in the treatment of diabetes mellitus, may also deserve attention with respect to possible effects on viral interactions with the its host cells, as recently pointed out (Scheen 2020, Lim *et al*. 2021).

It is striking that among eight genetic loci that affect the risk to contract severe COVID-19 when infected with SARS-CoV-2 (Pairo-Castineira *et al*. 2020), three carry allelic variants derived from Neandertals. This suggests that the genetic inheritance from Neandertals may have a larger impact on this disease than would naïvely be expected. However, local adaptation to infectious diseases often differs among human populations that have been separated by a few tens of thousands of years (Rees *et al.* 2020). Neandertals evolved largely independently from modern humans for about half a million years (Prüfer *et al.* 2014), even if rare genetic contacts occurred (Kuhlwilm *et al.* 2016; Meyer *et al.* 2016, Posth *et al*. 2017, Petr *et al.* 2020). Given the long time of separation, Neandertal adaptation to infectious diseases may therefore have differed drastically from that of modern humans.

The combination of large effect sizes and small number of Neanderal loci (and correspondingly smaller number of the multiple tests requiring correction) may allow associations with infection disease susceptibility to be detected in smaller cohorts than if all variants across the genome are considered. For the *DPP4* locus, we estimate that approximately two times more patients than currently available in HGI will be needed to achieve a 80% probability to detect the association between *DPP4* and severe COVID-19 with the standard genome-wide significance threshold (p<5e-8) (Supplementary Materials).

The three Neandertal genomes available to date, which vary in age between ~120,000 years and ~50,000 years and come from Europe and southern Siberia, are all homozygous for the risk variants on chromosome 2. Furthermore, the late Neandertal genome in Europe, which is most closely related to the Neandertals that mixed with modern humans, was homozygous also for the risk variants on chromosome 3 (Prüfer *et al*. 2017). It is thus likely that the risk variants at both these loci were at high frequency among late Neandertals. Barring other factors that may affect disease outcome, this means that, if alive today, a late Neandertal individual would have ~4-16 times higher risk of becoming critically ill if infected by SARS-CoV-2. This may support speculations that epidemic diseases could have played a role in the demise of Neandertals.

## Acknowledgments

We are indebted to the COVID-19 Host Genetics Initiative (HGI) for making the GWAS data available, to Tomislav Maricic for valuable input, and to the Max Planck Society and the NOMIS Foundation for funding.

## Supplementary Materials

### Neandertal gene-flow at the DPP4 locus

To detect introgression we first looked at alleles homozygously found in three high-coverage Neandertal genomes (Prüfer *et al*. 2014, Prüfer *et al*. 2017, Mafessoni *et al*. 2020) and absent in 108 Yoruba individuals, yielding two SNPs at *ACE2* and 39 and at *DPP4*. To further study the *DPP4* locus, we investigated all SNPs in linkage disequilibrium with rs117888248 (Supplementary Table S1) in the Neandertal genome (https://bioinf.eva.mpg.de/jbrowse/) most closely related to the Neandertal population that contributed genetically to present-day human (Prüfer *et al*. 2017). We demonstrated gene flow from Neandertals at the *DPP4* locus as previously described for the chromosome 3 locus (Zeberg and Pääbo, 2020a); using sites carrying alleles fixed among Neandertals and absent among Yoruba, phylogenetic trees, and the equation derived by Huerta-Sanchez *et al*. (2014). Previous genome-wide analyses have also found gene-flow at this locus (Vernot *et al*. 2014, Sankararam *et al*. 2012, Steinrücken *et al.* 2018, Chen *et al*. 2020, Skov *et al*. 2020).

### Introgression-wide significance

The three hitherto described Neandertal genetic risk factors for severe COVID-19 are all present at allele frequencies of ≥1% in European genomes from the 1000 Genomes Project and the COVID-19 HGI cohort. None of them are present in the three African populations in the 1000 Genomes Project with lowest amount of Neandertal admixture (Chen *et al*. 2020), *i.e*., Mende in Sierra Leone (MSL), Esan in Nigeria (ESN) and Yoruba in Ibadan, Nigeria (YRI). All three haplotypes carry derived alleles homozygously present in the three high-coverage Neandertal genomes (Prüfer *et al*. 2014, Prüfer *et al*. 2017, Mafessoni *et al*. 2020). We filtered SNPs fulfilling these criteria using a set of high-confidence Neandertal haplotypes defined by the intersection of previously published genome-wide maps of gene-flow from Neandertals (Vernot *et al*. 2014, Sankararam *et al*. 2012, Steinrücken *et al.* 2018, Chen *et al*. 2020, Skov *et al*. 2020). To exclude Neandertal-like variants due to incomplete lineage sorting, we further required the resulting haplotypes to have a length of at least 10 kb. Using these criteria, we identify 40,055 SNPs. We use these SNPs to estimate the effective number of hypotheses in European genomes from the 1000 Genomes Project using the Genetic Type I Error Calculator (Li *et al*. 2011). This yields a significance threshold of 8.7e-6 and a suggestive threshold of 1.7e-4, for a Neandertal “introgression-wide association study” (“IWAS”).

### Sample size needed to detect the DPP4 haplotype

As shown above, the power to detect a variant is improved if only introgressed Neandertal haplotypes are considered, although under different null-hypotheses. We calculated the sample size needed to achieve genome-wide significance (p<5e-8), using standard techniques (Pirinen *et al*. 2020). If there is a non-zero effect, *i.e.*, β≠0, then the *z*-score is distributed as *z◻N*(β/SE,1) and *z*^2^◻χ^2^_1_((β/SE)^2^). The parameter (β/SE)^2^ is known as the *non-centrality parameter* and scales linearly with sample size. The non-central chi-squared distribution was used to calculated the probability of observing a sufficiently strong association, *i.e.* statistical power. To reach 80% power to detect the Neandertal *DPP4* haplotype we find that we need approximately twice the sample size. For 99% detection probability the sample size needs to tripple.

## Data availability

The archaic genomes are availability at the server of the Max Planck Institute for Evolutionary Anthropology (http://cdna.eva.mpg.de/neandertal/Vindija/VCF/ and https://bioinf.eva.mpg.de/jbrowse/) and the modern human genomes at the 1000 Genomes Project server (http://ftp.1000genomes.ebi.ac.uk/vol1/ftp/release/20130502/). GWAS summary statistics can be obtained from the COVID-19 Host Genetics Imitative (https://www.covid19hg.org/results/, round 4 release: A/B2_ALL_eur_leave_23andme).

## Supplementary Table

**Supplementary Table S1.** SNPs in linkage disequilibrium with rs117888248 and the corresponding Neandertal alleles. LD data from the 1000 Genomes Project, “Vindija” refers to a Croatian Neandertal genome (https://bioinf.eva.mpg.de/jbrowse/).

| Chr | Pos | rsid | LD with rs117888248 | Ref | Alt/Risk | Vindija |
| --- | --- | --- | --- | --- | --- | --- |
| 2 | 162936216 | rs117888248 | 1.000 | C | T | T/T |
| 2 | 162938711 | rs79624636 | 1.000 | G | C | G/C |
| 2 | 162933386 | rs116906287 | 1.000 | A | G | A/G |
| 2 | 162932814 | rs118098838 | 1.000 | G | T | T/T |
| 2 | 162940017 | rs76135328 | 1.000 | C | A | A/A |
| 2 | 162941569 | rs117460501 | 1.000 | A | G | G/G |
| 2 | 162944795 | rs117965144 | 1.000 | C | T | T/T |
| 2 | 162945153 | rs117516594 | 1.000 | G | A | G/A |
| 2 | 162949980 | rs188765526 | 1.000 | G | A | G/G |
| 2 | 162955612 | rs145259049 | 1.000 | C | T | C/T |
| 2 | 162956598 | rrs77914221 | 1.000 | T | C | T/C |
| 2 | 162957351 | rs57355997 | 1.000 | T | A | A/A |
| 2 | 162938141 | rs75778040 | 0.908 | A | T | T/T |
| 2 | 162932254 | rs6709208 | 0.899 | C | T | T/T |
| 2 | 162931074 | rs115498481 | 0.808 | G | T | T/T |

## Notes

### Competing Interest Statement

The authors have declared no competing interest.

