## Supplementary figures and images for "The MERS-CoV receptor gene is among COVID-19 risk factors inherited from Neandertals"

### Supplementary Table S1

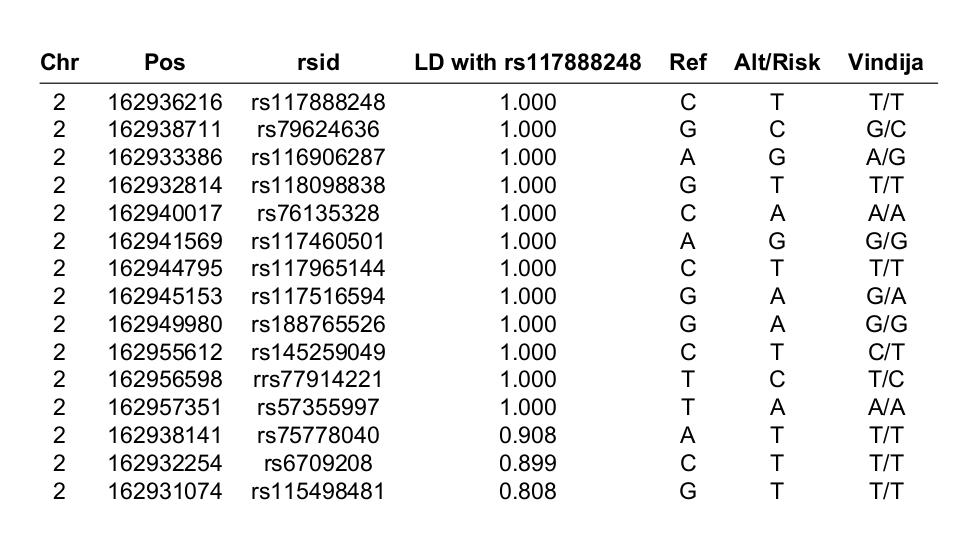
